# Longitudinal single-cell immune profiling revealed distinct innate immune response in asymptomatic COVID-19 patients

**DOI:** 10.1101/2020.09.02.276865

**Authors:** Xiang-Na Zhao, Yue You, Guo-Lin Wang, Hui-Xia Gao, Xiao-Ming Cui, Li-Jun Duan, Sheng-Bo Zhang, Yu-Ling Wang, Lin-Yao, Li Li, Jian-Hua Lu, Hai-Bin Wang, Jing-Fang Fan, Huan-Wei Zheng, Er-Hei Dai, Lu-Yi Tian, Mai-Juan Ma

## Abstract

Recent studies have characterized the single-cell immune landscape of host immune response of coronavirus disease 2019 (COVID-19), specifically focus on the severe condition. However, the immune response in mild or even asymptomatic patients remains unclear. Here, we performed longitudinal single-cell transcriptome sequencing and T cell/B cell receptor sequencing on 3 healthy donors and 10 COVID-19 patients with asymptomatic, moderate, and severe conditions. We found asymptomatic patients displayed distinct innate immune responses, including increased CD56^bri^CD16^−^ NK subset, which was nearly missing in severe condition and enrichment of a new Th2-like cell type/state expressing a ciliated cell marker. Unlike that in moderate condition, asymptomatic patients lacked clonal expansion of effector CD8^+^ T cells but had a robust effector CD4^+^ T cell clonal expansion, coincide with previously detected SARS-CoV-2-reactive CD4^+^ T cells in unexposed individuals. Moreover, NK and effector T cells in asymptomatic patients have upregulated cytokine related genes, such as *IFNG* and *XCL2*. Our data suggest early innate immune response and type I immunity may contribute to the asymptomatic phenotype in COVID-19 disease, which could in turn deepen our understanding of severe COVID-19 and guide early prediction and therapeutics.

## INTRODUCTION

Severe acute respiratory syndrome coronavirus 2 (SARS-CoV-2), the causative agent of the coronavirus disease 2019 (COVID-19), has rapidly caused a worldwide pandemic with ever-increasing cases and COVID-19-related deaths (Organization, 2020). COVID-19 patients exhibit a broad spectrum of clinical manifestation, ranging from mild or even asymptomatic infection to severe disease or death (Raoult et al., 2020). Therefore, understanding the host immune response involved in the disease course is of supreme importance to the development of effective therapies.

In severe COVID-19 patients, hyper-inflammation responses referred to as a cytokine storm (Jose and Manuel, 2020; Mehta et al., 2020) and lymphopenia (Chen et al., 2020a; Huang et al., 2020) have been considered risk factors associated with the detrimental progression of COVID-19 patients. Elevated pro-inflammatory cytokines (e.g., IL-1β, IL-6, and TNF-α) and inflammatory monocytes and neutrophils and a sharp decrease in lymphocytes have also been reported in severe patients (Chen et al., 2020a; Chen et al., 2020b; Giamarellos-Bourboulis et al., 2020; Huang et al., 2020; Jose and Manuel, 2020; Lucas et al., 2020; Mathew et al., 2020; Skarica et al., 2011; Zhou et al., 2020). Further, single-cell RNA sequencing (scRNA-seq) studies of peripheral blood mononuclear cells (PBMCs) or bronchoalveolar lavages have reported the immunological profiles to SARS-CoV-2 in severe and moderate patients (Lee et al., 2020; Liao et al., 2020; Mathew et al., 2020; Wilk et al., 2020; Zhang et al., 2020; Zhu et al., 2020), suggesting moderate disease was associated with more protective T cell-dependent response, with exacerbated systemic inflammation and less effect T cells in severe disease. Longitudinal immune responses of moderate and severe COVID-19 patients have been analyzed by flow cytometry (Lucas et al., 2020), while unbiased longitudinal single-cell transcriptome profiling is still missing. On the other hand, the contribution of asymptomatic individuals to the transmission of SARS-CoV-2 rise a significant public health concern (Gandhi et al., 2020). Despite the clinical and immunological assessment of asymptomatic individuals (Long et al., 2020), transcriptome profiles of asymptomatic individuals is lacking, which might help us understand the nature of the asymptomatic COVID-19 disease.

To elucidate characteristics that might lead to immunopathology or protective immunity in asymptomatic, moderate, and severe COVID-19, we performed scRNA-seq together with single-cell V(D)J sequencing to longitudinal analyses the immunological profile of PBMCs from ten hospitalized COVID-19 patients and three healthy donors, and to identify correlations between distinct immune phenotype and disease severity.

## RESULTS

### Single-cell transcriptomes profiling of PBMCs from COVID-19 patients

In order to explore the immunological changes of patients with COVID-19, the immune profiles of PBMCs from ten patients and three healthy donors (HD) were analyzed by scRNA-seq with T cell and B cell receptor (TCR/BCR) sequencing using the 10x Chromium platform (**Figure 1a**). The demographics and clinical features of these patients are shown in **Table S1**. The ten patients were classified into severe (Ps, n=1), moderate (Pm, n=7), and asymptomatic (Pa, n=2), aged 17 to 62 years old, and 6 of them were male. A total of 31 blood samples were collected from 10 patients, and 8 of 10 patients provided blood samples ≥ three times at different time points during hospitalization (**Figure 1a**). For moderate and severe patients, the days were recorded based on the time after their symptom onset. The days were recorded for asymptomatic patients based on the time they confirmed SARS-CoV-2 positive by RT-PCR test.

**Figure 1.**
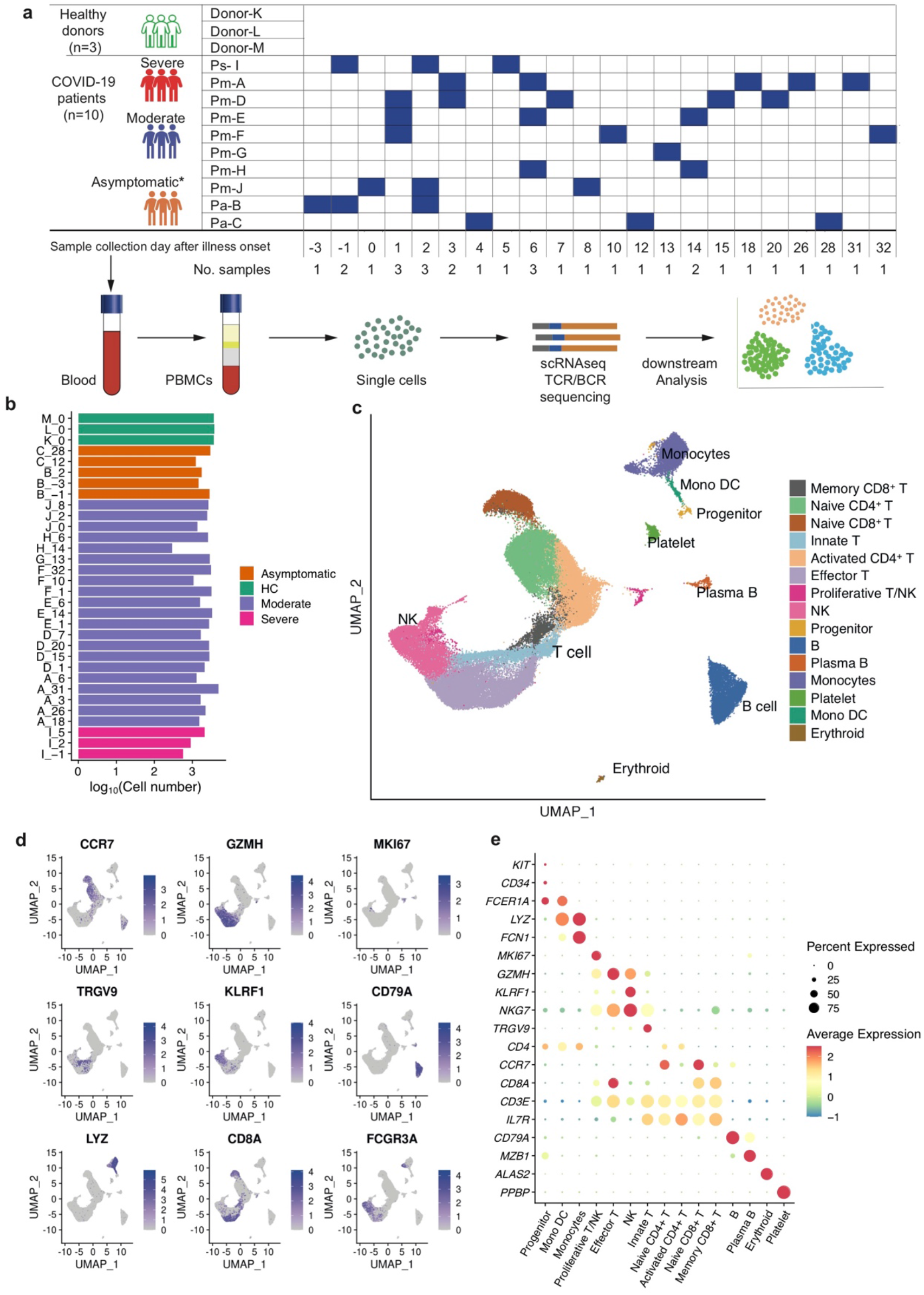
Single-cell RNA sequencing of peripheral blood cells from COVID-19 patients. **a**, Timeline of blood sample collection in the 13 subjects (1 severe, 7 moderate, 2 asymptomatic, and 3 healthy donors) and schematic outline of the study design. **b,** Bar plot shows the log10 transformed cell number of each sample for each subject at different time points. Green, orange, blue, and red represent samples collected from healthy donors, asymptomatic, moderate, and severe patients, respectively. The same color palette was used throughout the study. **c,** UMAP of cells after data integration. Cell types were identified by the marker genes after clustering. Each dot represents an individual cell. A total of 15 cell types were identified and color-coded. **d,** Canonical cell markers that used to annotate clusters as represented in the UMAP plot. Colored according to expression levels and legend labeled in log scale. **e**, Dot plots of average expression and percentage of expressed cells of marker genes in each labeled cell type.

After removing low-quality cells and doublets, we kept 73592 cells for downstream analysis, including 62386 cells from ten patients and 11206 cells from three healthy donors. On average, there were 2300 cells for each participant at each time point (**Figure 1b**). We identified 15 major cell types after data integration and unsupervised clustering (**Figure 1c-e; Figure S1a, 1c**), including innate T cells (*TRGV9*^+^), effector T cells (*GZMK*^+^), naive CD8^+^ T cells (*CCR7*^+^*SELL*^+^), memory CD8+ T cells (*GPR183*^+^), naive CD4^+^ T cells (*CCR7*^+^ *SELL*^+^), activated CD4^+^ T cells (*IL7R*^+^*CCR7*^−^), proliferative T/natural killer (NK) cells (*MKI67*^+^), NK cells (*NKG7*^+^), progenitor cells (*CD34*^+^GATA2^+^), B cells (*CD79A*^+^*MS4A1*^+^), plasma B cells (*CD38*^+^*MZB1* ^+^), monocytes (CD14^+^ monocytes: *LYZ* ^+^; CD16^+^ monocytes: *FCGR3A*^+^), platelet (*PPBP*^+^), monocyte-derived dendritic cells (Mono DC: *CD1C*^+^), and erythroid cells (*ALAS2*^+^).

To quantify changes in major cell types, we investigated the relative proportions of 15 cell types among PBMCs in different disease conditions and stages (**Figure 2a-c; Figure S1a-c**). Unlike asymptomatic and moderate patients, severe patients had significant changes across multiple cell types, especially for a significantly increased proportion of NK cells (**Figure 2a**). Further analysis with unsupervised clustering on relative proportions of the 15 major cell types from all samples revealed a marked increase of NK cells, plasma B cells and platelets, and a decrease of CD4^+^ and CD8^+^ T cells in severe patients (**Figure 2b**). We summarised the samples from moderate patients into 3 stages, based on the days after symptom onset (<10 days, 10-20 days, and >20 days), and this classification was used throughout the study. Further statistical analyses showed that the proportions of T cell subsets were highly heterogeneous among different stages in moderate and asymptomatic patients, including activated CD4^+^ T cells and memory CD8^+^ T cells with consistently lower abundance in severe and healthy conditions (**Figure 2c; Figure S2a, 2b**). In contrast, the relative proportion of NK cells, plasma B cells, and platelets was increased in severe patients (**Figure 2c**).

**Figure 2.**
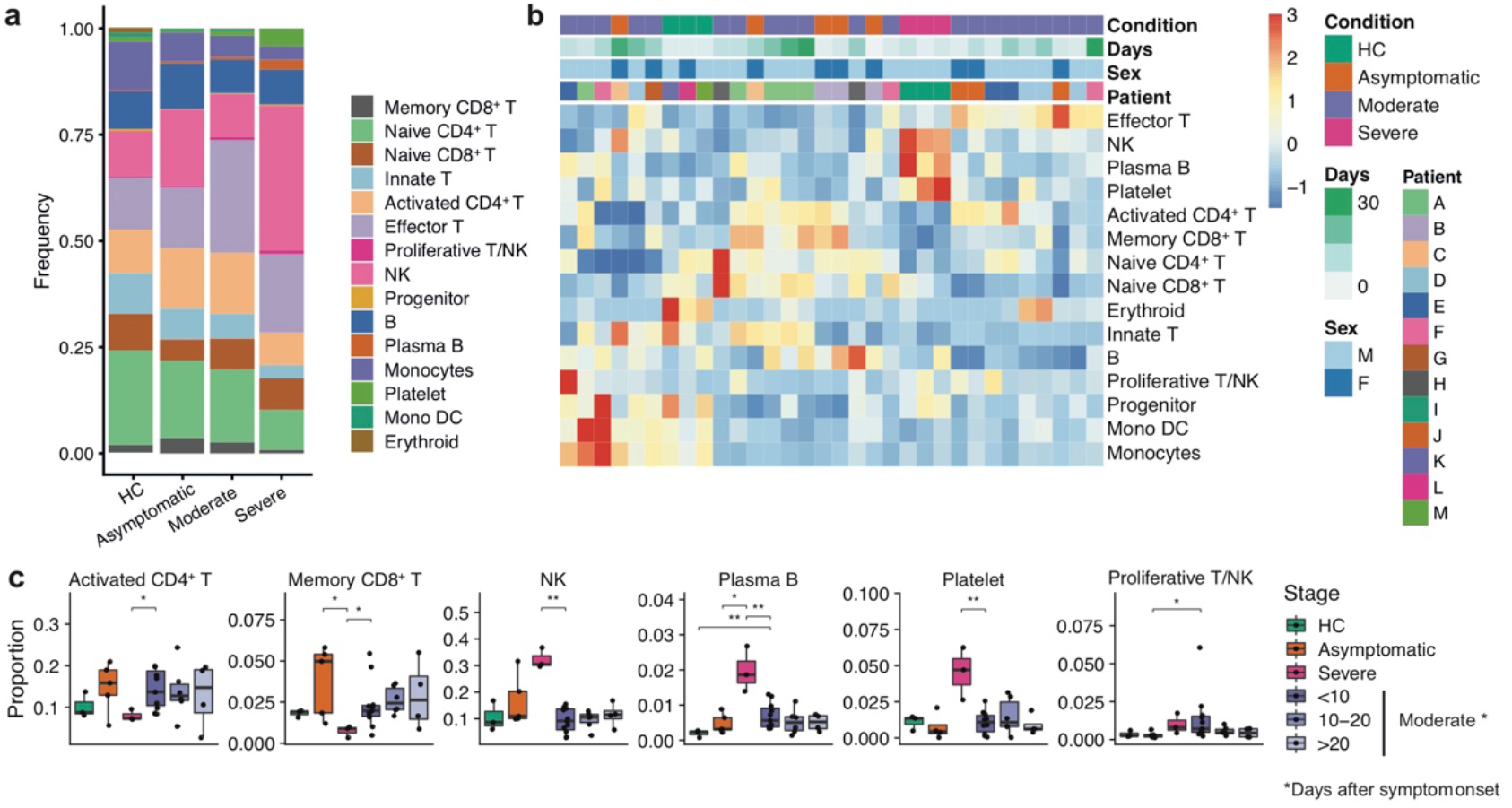
Differences in major cell types compositions across disease conditions. **a,** Proportion of cell types in PBMCs of healthy donors (n=3), moderate (n=7), severe (n=1), and asymptomatic patients (n=2). Colored according to cell type information. **b**, Heatmap of 15 cell types identified in **Figure 1c** according to disease conditions, days after illness onset, and sex. The color scale is based on row-scaled z-score. **c**, Boxplots showing the percentages of each cell type in four disease conditions (HD, asymptomatic, moderate, and severe) and stages. Moderate patient samples were divided into three stages based on days after symptom onset. Boxes are colored according to disease conditions and stages of the moderate condition. Two-sided unpaired Mann–Whitney U-test was used for analysis, and a *p*-value < 0.05 is considered significant. \**p*<0.05, \*\**p*<0.01, \*\*\**p*<0.001.

### Immune profiles of T cells and NK cells in COVID-19 patients

To further characterize the T/NK subsets in detail, we extracted the data from T/NK cells and repeated the data integration and clustering. With the refined clustering analysis, we identified 14 cell subtypes (**Figure 3a, b**), including rare clusters absent in previous clustering analysis (**Figure 1c; Figure S3a**). We identified four CD4^+^ T cell subsets, including naive CD4^+^ T cells (T_N_, *CCR7*^+^), CD4^+^ central memory T cells (T_CM_, *GPR183*^+^*CCR7*^+^), CD4^+^ effector memory T cells (T_EM_, *CCR7*^−^*SELL*^−^*GZMA*^+^), and CD4^+^ effector cells (T_E_, *GZMA*^+^*GZMB*^+^). We identified three CD8^+^ T cell subsets, including CD8^+^ T_N_ cells (*CCR7*^+^), CD8^+^ T_E_ cells (*GZMA*^+^*GZMB*^+^), and CD8^+^ T_CM_ cells (*GPR183*^+^). Additionally, we identified seven innate immune subsets, including mucosal-associated invariant T cells (MAIT, *SLC4A10*^+^*TRAV1-2*^+^), gamma-delta T (γδT) cells (*TRGV9*^+^*TRDV2*^+^), immature NK cells (iNK: *KIT*^+^), CD56^bri^CD16^−^ NK cells, CD56^dim^CD16^+^ NK cells, and a previously uncharacterized Th2-like lymphoid population (*CD4*^−^*CD8A*^−^*PTGDR2*^+^). We also identified a proliferative T/NK population (*MKI*67^+^).

**Figure 3.**
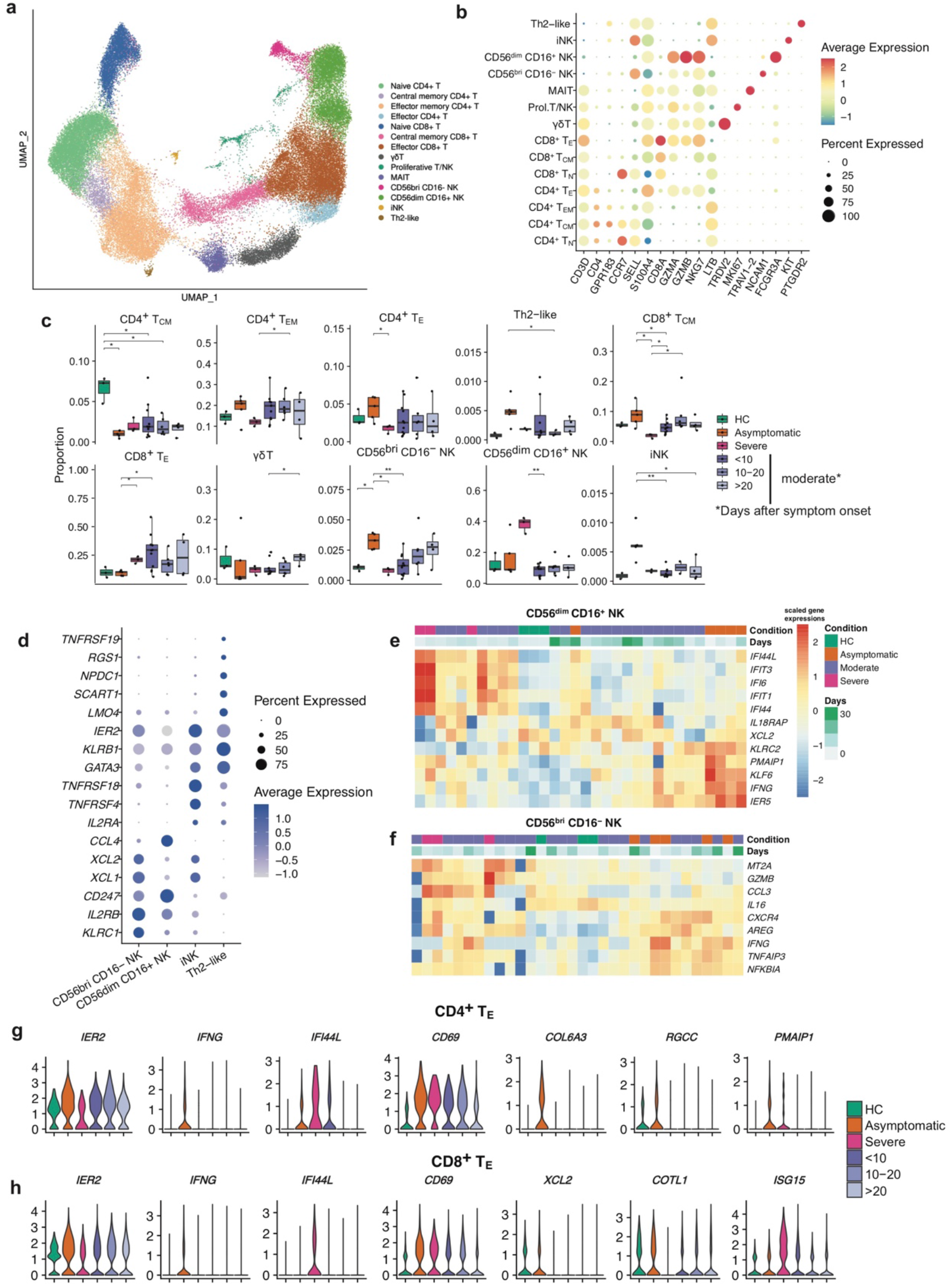
Identification and characterization of the subpopulation of T and innate immune cells in COVID-19 patients. **a**, UMAP of T and NK cells by Seurat. Cell types were identified by the marker genes. Each dot represents an individual cell. A total of 7 T cell subtypes and 7 innate immune cell subtypes were identified and color-coded. **b**, Dot plot of average expression and percentage of expressed cells of selected canonical markers in each labeled cell subtype. **c**, Proportions of subtypes in each sample in different disease conditions. Two-sided unpaired Mann– Whitney U-test was used for analysis, and a *p*-value < 0.05 is considered significant. **d**, Dot plot showing the average expression and percentage of expressed cells of selected differentially expressed genes (DEGs) between 3 NK subtypes and Th2-like subset. **e-f**, Heatmap of selected DEGs between different condition for CD56^bri^CD16^−^ (**e**) and CD56^bri^CD16^+^ (**f**) NK cell subtypes. The color scale is based on row-scaled z-score of gene expression. **g-h**, Violin plots showing expression of selected DEGs in CD4^+^ (**g**) and CD8^+^ (**h**) effector T cell subsets. \**p*<0.05, \*\**p*<0.01, \*\*\**p*<0.001.

To gain insights into features in T/NK cell subtypes, we assessed the abundance of each cell type across disease conditions and stages (**Figure 3c; Figure S4b**). Notably, the proportion of CD4^+^ T_E_ significantly increased in asymptomatic patients compared with severe patients, while the moderate and severe patients had increased CD8^+^ T_E_ (**Figure 3c**). We also observed an increased proportion of CD8^+^ T_cm_ in asymptomatic patients compared with other disease conditions and stages, and all patients had a decreased proportion of CD4^+^ T_cm_ compared to healthy donors (**Figure 3c**). Of interest, the abundance of CD56^bri^CD16^−^ NK was significantly higher in asymptomatic patients than other conditions and was increased in moderate patients over time (**Figure 3c**). In contrast, the CD56^dim^CD16^+^ NK, which was the most abundant NK subset, was substantially enriched in severe patients. Like CD56^bri^CD16^−^ NK, its precursor iNK also increased in asymptomatic patients. In addition, we found that the Th2-like lymphoid had increased abundance in asymptomatic patient samples and some early moderate patient samples (<10 days post symptom onset). These results indicated that asymptomatic patients had distinct innate immune cell distribution compared with other conditions.

Next, we sought to identify the specific signature of the NK and Th2-like innate immune cells with distinct distribution in asymptomatic and severe conditions. We found that CD56^bri^CD16^−^ NK cells have high expression of *XCL1*, *XCL2*, and *IFNG* (**Figure 3d**), consistent with our knowledge that these cells are efficient cytokine producers (Michel et al., 2016). The Th2-like lymphoid cells were *TCR^−^CD3^−^CD4^−^CD8^−^* but expressed Th2 markers such as *PTGDR2* and *GATA3*, they were classified mainly as Th2-like cells by singleR (**Figure S5a**). In addition, Th2-like cells expressed myeloid cell markers such as *LMO4*, which is also expressed in eosinophils/basophils/mast cells (Herman et al., 2018), while these cell types were not present in this dataset. We also found that *TNFRSF19* uniquely expressed in the Th2-like cells in this dataset (**Figure S5b**). Interestingly, although *TNFRSF19* is absent in most of the immune cells according to previous study (Kojima et al., 2000) and the human lung cell atlas database (Travaglini et al., 2020). It is highly expressed in epithelial cells such as ciliated cells, which also express ACE2 and are considered as entry cells of SARS-CoV-2 (**Figure S5c**). The Th2-like cells also expressed *NPDC1* and *RGS1*, which has been linked to human tissue-resident memory T cells and gut T cell trafficking correspondingly (Gibbons et al., 2011; Kumar et al., 2017).

To further investigate the difference of transcriptomes for each cell type of T and NK cells across different conditions, we performed systematic differential gene expression (DGE) analysis. We found distinct signatures expressed in severe and asymptomatic patient samples in NK cells and effector T cells (**Figure S6a-e**). Genes related to type I interferon (IFN-I) signaling pathway was widely upregulated in severe conditions (**Figure S6b**). We observed upregulated chemokine *XCL2* and *CXCR4* and cytokine IFNG in CD56^dim^CD16^+^ NK and CD56^bri^CD16^−^ NK cells in asymptomatic conditions (**Figure 3e-3f; Figure S6a, 6c**). Effector CD4^+^ and CD8^+^ T cells in asymptomatic conditions were marked by the upregulation of distinct genes, including immediate early genes (*IER2* and *IER5*) and *IFNG* (**Figure S6d-e**). We observed an upregulated expression of *COL6A3* in CD4^+^ effector T cells and NK cells of asymptomatic conditions. *COL6A3* encodes type VI collagen, and the expression of collagens in structure cells is expected to enhance their ability to interact with hematopoietic cells(Krausgruber et al., 2020). However, the expression of collagens in immune cells is less discussed, and we suspect they might also facilitate better interaction with structure cells.

### Clonal expansion in T cells and usage of V(D)J genes COVID-19 patients

To evaluate the clonal relationship among individual T cells and usage of V(D)J genes across different conditions, we also analyzed single-cell TCR sequencing data and reconstructed high-quality TCR sequence in 70.5% of the T cells with various degrees of clonal expansion among T cell subsets (**Figure 4a; Figure S7a-d**). The clonal expansion was evident in patients compared with healthy donors, whereas moderate patients presented higher clonal expansion than those of the asymptomatic and severe conditions (**Figure 4b-d**). In addition, clonal expansion was enriched in CD4^+^ and CD8^+^ effector T cell in moderate patients, whereas CD4^+^ effector T cells in asymptomatic patients have more clonal expansion (**Figure 4c**), which was consistent with the distribution of T cell subsets that asymptomatic patients have more CD4^+^ effector T cells instead of CD8^+^ effector T cells. Of note, samples from moderate conditions had high clonal expansion and low diversity at the early stage and decreased over time, indicating a recovery of the disease (**Figure 4d**). Meanwhile, large clonal expansions (clonal size >200) were absent in the severe condition (**Figure 4b**), indicating that severe patients might lack efficient clonal expansion of effector T cells during early-stage infection of SARS-CoV-2.

**Figure 4.**
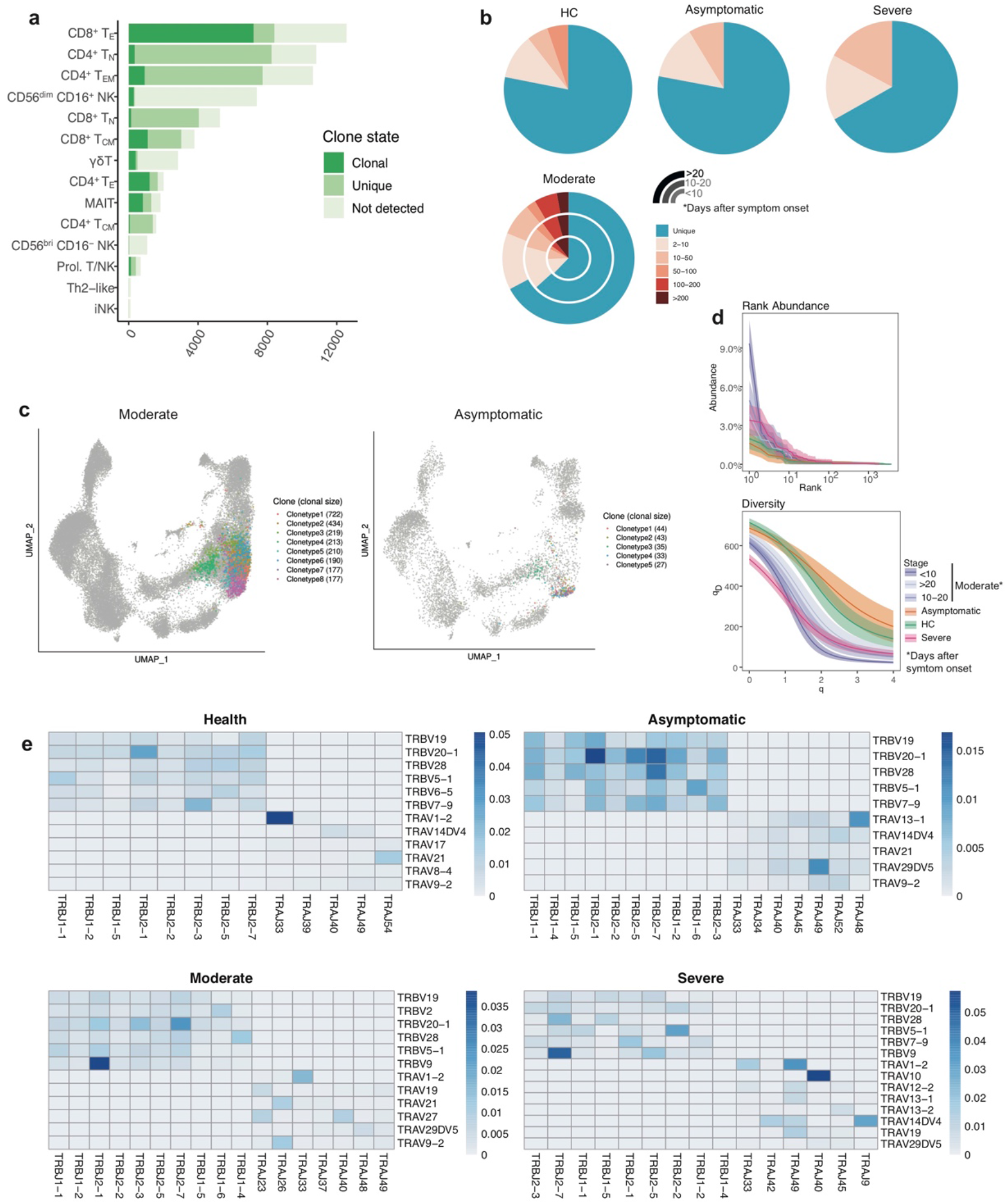
Expanded TCR clones and selective usage of V(D)J genes. **a,** Bar plots showing the numbers of TCR detected in each T and NK cell subtype. **b**, The percentage of the clonal status of T cells with TCR. The clonal status was defined by clone size in four disease conditions. **c**, UMAP of T cells derived from PBMCs for moderate and asymptomatic patients. Clusters are denoted by colors labeled with TCR clones with top5 largest clone size in moderate patients, and TCR clones with top5 largest clone size are shown in asymptomatic patients. **d**, TCR abundance and diversity across disease conditions and stages generated by alakazam package. The 95% confidence interval is estimated via bootstrapping (B = 200). **e**, Heatmaps of the difference in TRA/B rearrangement in four conditions. The colors represent the percentage of V-J gene usage.

We compared the usage of V(D)J genes across disease conditions and disease stages of moderate patients (**Figure 4d-e; Figure S7e-f**). We observed a different usage of V(D)J genes with decreased diversity in severe patients, which was more pronounced in TRA genes (**Figure 4e**). We also found that over-representation of *TRAJ40* and *TRBJ2-7* in severe patients compared to moderate and asymptomatic patients (**Figure 4e; Figure S7e-f**). The preferred TRBJ gene in severe patients was *TRBJ40*, whereas *TRBJ2-1* was preferred in moderate and asymptomatic patients (**Figure 4e**). The selective usage of V(D)J genes indicated that different immunodominant epitopes may drive the molecular composition of T cell responses and may be associated with SARS-CoV- 2-specific infection.

### Features of B cells and expansion and specific rearrangements of V(D)J genes

We extracted single-cell B cells sequencing data and repeated the data integration and clustering analysis. We identified three B cell subsets according to the expression of canonical B cell markers, including naïve B cells (*MS4A1*^+^*IGHD*^+^), memory B cells (*CD27*^+^), and plasma B cells (*MZB1*^+^*CD38*^+^) (**Figure 5a, b**). Severe patients and early-stage samples (<10 days after symptom onset) from moderate patients had a higher proportion of plasma B, and a declining trend of the plasma B proportion was observed over time for the samples of moderate patients (**Figure 5c**).

**Figure 5.**
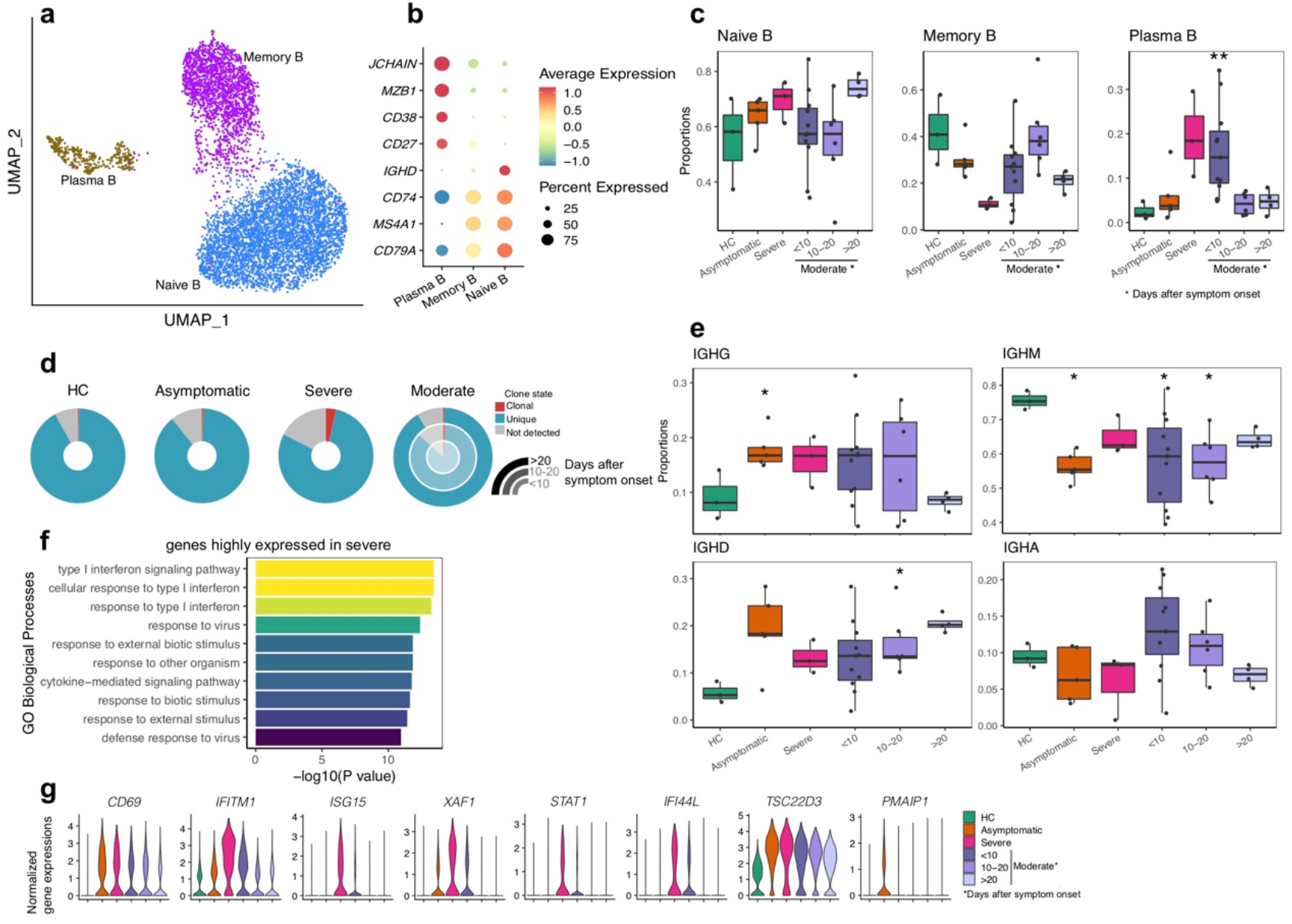
Subpopulation analysis of B cells. **a**, UMAP projection of B cells. Each dot represents a single cell, colored according to cell type. **b**, Dot plot of canonical cell markers used to annotate clusters in the UMAP plot. **c**, Boxplots showing the differences in percentages of each cell type comparing different disease conditions to healthy control. Two-sided unpaired Mann–Whitney U-test was used for analysis, and a *p*-value < 0.05 is considered significant. **d**, The percentage of clonal status in B cells that have BCR, across disease conditions and stages. **e**, Boxplots showing the proportion of IgG, IgM, IgD, and IgA comparing different disease conditions to healthy control. Two-sided unpaired Mann–Whitney U-test was used for analysis, and a *p*-value < 0.05 is considered significant. **f**, Bar plot of top 10 enriched GO terms generated from genes upregulated in severe conditions. **g,** Violin plots of selected DEGs generated by comparing different conditions. \**p*<0.05, \*\**p*<0.01, \*\*\**p*<0.001.

Next, we analyzed the single-cell BCR sequencing data and reconstructed high-quality BCR sequences in more than 80% of the B cells. We found an apparent BCR clonal expansion in severe patients compared to asymptomatic, moderate, and healthy donors (**Figure 5d**), indicating robust activation of B cell activity and humoral immune responses in severe patients. There was less clonal expansion in moderate patients, and the clonal expansion decreased over time (**Figure 5d**), suggesting humoral immune responses reduced at the convalescent stage. We also evaluated the distribution of IgG, IgM, IgD, and IgA at different disease conditions and stages (**Figure 5e**). Overall, all patients had higher IgG and lower IgM compared to healthy donors, especially for asymptomatic patients, who also had significantly higher IgG and low IgM (**Figure 5e**). The IgG and IgM were highly variable at the early moderate patient sample and returned to similar levels of healthy donors at the late stage (>20 days post symptom onset).

Next, we assessed V(D)J rearrangements of the BCR and analyzed the usage of V(D)J genes across different disease conditions and stages (**Figure S8a**). We found more specific V(D)J usage in the severe condition than the other three conditions, indicating that B cells might have undergone unique and specific V(D)J rearrangements. We also identified comprehensive usage of *IGHJ4* and *IGLJ3* in all patients, but the paired *IGHV* genes of *IGHJ4* and *IGLJ3* were different in severe patients (*IGHV*3*−33* and *IGLV1−51*, respectively) compared with asymptomatic and moderate conditions (*IGHV3−23* and *IGLV1−44*, respectively) (**Figure S8a**). Moreover, the top two paired V-J frequencies in severe patients were *IGLV1−51/IGKJ3* and *IGHV3−33/IGHJ4*, whereas IGHV3−23/IGHJ4 and *IGLV1−44/IGLJ3* for asymptomatic patients, and *IGLV1−44/IGLJ3* and *IGLV2−14/IGLJ2* for moderate patients (**Figure S8a**).

To further investigate transcriptomic changes in B cells after SARS-CoV-2 infection, we performed a DGE analysis comparing different conditions in each B cell subsets (**Figure 5f**). We found increased expression of *CD69* and *TSC22D3* in disease compared to healthy controls. Like T/NK cells, we found an increased expression of marker genes, including *IFITM1*, *ISG15*, *XAF1*, and *IFI44L*, involving the IFN-I signaling pathway in the severe and early stage of moderate patients (expression decreased over time) (**Figure 5g**). However, severe patients have much higher expression of these genes (**Figure 5g**), indicating more robust activation of IFN-I signaling pathway response and a pro-inflammatory state. Notably, we also found increased expression of *STAT1* in severe patients (**Figure 5g**), which was consistent with a previous study (Zhu et al., 2020).

## DISCUSSION

In this study, we performed longitudinal scRNA-seq and V(D)J-sequencing on PBMCs from COVID-19 patients in different conditions from asymptomatic to severe. A part of the result generated in this study was consistent with recent studies. First, we confirmed that early-stage severe patients had increased type I interferon (IFN-I) response in most of the immune cells compared to other conditions, which might contribute to a hyper-inflammatory response. The longitudinal analysis revealed that the early-stage moderate patients also showed increased IFN-I response but less than severe condition, and the response declined with time. While we only have samples from the early stage in a severe patient and cannot assess the change in late stage, another study has shown that the cytokines related to IFN-I, such as IFN-α, did not decrease in severe condition over time (Lucas et al., 2020). Second, we found different proportions of immune subsets in different conditions that consistent with recent studies (Lee et al., 2020; Liao et al., 2020; Lucas et al., 2020; Wilk et al., 2020; Zhang et al., 2020; Zhu et al., 2020), including the increased plasma B cells and decreased T subsets in severe conditions. Higher proportions of T subsets were marked in moderate condition, and TCR analysis showed increased clonal expansion of T cells compared to other conditions (Zhang et al., 2020). The BCR analysis revealed increased clonality of B cells in severe conditions, with decreased IgM and increased IgG proportions in all disease conditions compared to healthy controls, which came back to normal levels in late-stage moderate patients. We did not observe a significant increase of monocytes in severe conditions (Schulte-Schrepping et al., 2020), which might due to the sampling time that only early-stage severe samples were included in this study. Collectively, the severe condition was marked by a hyper-inflammatory immune response, including upregulated IFN-I response, impaired T cell clonal expansions, and increased plasma B cells and BCR clonality. In contrast, the moderate patients have activated, but relatively lower expression of IFN-I related genes and robust T cell clonal expansions at early-stage with decreased immune response at later-stage, indicating convalescence.

Although different immune responses between moderate and severe conditions have been the focus of many recent studies (Lee et al., 2020; Liao et al., 2020; Wilk et al., 2020; Zhang et al., 2020; Zhu et al., 2020), the mechanisms under asymptomatic phenotype of COVID-19 are less studied and have become the elephant in the room. In this study, we performed comprehensive comparisons between asymptomatic and other conditions, and found that asymptomatic condition is not just an intermediate state between healthy and moderate but has its unique immunological features. PBMCs in asymptomatic patients had lower IFN-I related gene expression than in severe and early-stage moderate conditions. Asymptomatic patients did not show strong clonal expansion of BCR and TCR, but they had a robust effector CD4^+^ T cells clonal expansion and less effector CD8^+^ T cells expansions, while the former was lacking in the severe patient. Moreover, the CD4^+^ effector T cells in asymptomatic patients had upregulated *IFNG* compared to other conditions, which together with the increased CD4^+^ T cells suggested an activated type 1 immunity. CD4^+^ effector T cells play an essential role in regulating the antiviral inflammatory response and mediating viral clearance through direct cytotoxic effects on virus-infected cells (Swain et al., 2012), it could also recognize SARS-CoV-2 by cross-reactive epitopes in unexposed individuals (Grifoni et al., 2020). We hypothesize that the increased activity of effector CD4^+^ T cells in asymptomatic patients may originated from pre-existing immunity to SARS-CoV-2 and promote early rapid control of the disease.

Despite the difference in adaptive immune response, there were profound differences in innate immune responses between asymptomatic condition and others. Asymptomatic patients had significantly increased fractions of CD56^bri^CD16^−^ NK cells and their precursor (iNK), while the severe patient had increased CD56^dim^CD16^+^ NK subsets. Moderate patients had fewer CD56^bri^CD16^−^ NK cells at the early stage, and the fraction was increased at the later stage. We found CD56^bri^CD16^−^ NK subsets had upregulated cytokine-related genes such as *XCL1*, *XCL2*, and *IFNG*, consistent with current knowledge that these regulatory cells act as potent cytokine and chemokine producers. CD56^dim^CD16^+^ NK cells are considered to be differentiated from CD56^bri^CD16^−^ NK cells and have more cytotoxic potential (Abel et al., 2018). In addition to the difference in abundance, there were significant differences in gene expressions between severe and asymptomatic conditions, with cytokine related genes such as *IFNG* and *XCL2* upregulated in asymptomatic conditions. CD56^bri^CD16^−^ NK cells have been linked to virus infection. Infection of influenza A Virus induced NK cell hyperresponsiveness and cytokine production, particularly in the CD56^bri^CD16^−^ NK subset (Scharenberg et al., 2019). An asymptomatic hemophiliac patient coinfected with HIV/HCV also had increased CD56^bri^CD16^−^ NK cells (Fregni et al., 2013). These results suggested that the CD56^bri^CD16^−^ regulatory NK cells may play a critical role in the protection of SARS-CoV-2 infection in asymptomatic patients.

On top of the NK cells, we also identified a previously uncharacterized Th2-like innate immune cell subset enriched in asymptomatic patients. Interestingly, the Th2-like cells uniquely expressed gene *TNFRSF19*, which is rarely expressed in immune cells (Kojima et al., 2000). However, it is highly expressed in epithelial cells, which also express ACE2 and was considered as entry cells of SARS-CoV-2 infection (Sungnak et al., 2020). These cells also expressed genes related to tissue-homing and immune cell migration. Although the role of these Th2-like lymphoid cells in response to SARS-CoV-2 infection is unclear, we speculate that these cells might interact with infected epithelial cells.

There are several limitations to this study. Our sample size is relatively small,with only one severe and two asymptomatic patients. In addition, only early-stage blood samples were available from the severe patient. Therefore, future studies with longitudinal samples from more COVID-19 patients, especially for asymptomatic patients, are needed to further identify the relationships between immune characteristic and asymptomatic infection. Second, only blood samples from patients were used for immunological analyses, but not bronchoalveolar lavage fluid samples because of the difficulty of obtaining these samples due to biosafety reasons.

Overall, our analyses provide the first comprehensive characterization of the immune landscape of asymptomatic COVID-19 patients and highlight the importance of innate immune response towards disease progression. The reduced abundance of CD56^bri^CD16^−^ NK subset in the early-stage severe patient suggested avenues for both diagnostic and clinical intervention at the early stages of the disease, considering drugs such as Natalizumab could induce the expansion of CD56^bri^CD16^−^ NK cells (Skarica et al., 2011).

## Acknowledgments

We thank all patients for their participation in this study and for proving blood samples. We also thank Matt Ritchie, Peter Hickey, Gordon Smyth, and Yifan Zhan for their helpful discussions. This work was supported by grants from the Natural Science Foundation of China (81773494 to M-J.M.), the National Major Project for Control and Prevention of Infectious Disease of China (2017ZX10303401-006 to M-J.M.), the Special National Project on Investigation of Basic Resources of China (2019FY101502 to M-J.M.), and Emergency Science and Technology Project for Prevention and Control of COVID-19 (20277734D to E-H.D).

## Author Contributions

MJM, LYT, and EHD conceived and designed the study. HXG, YLW, LL, JHL, HBW, JFF, and HWZ collected clinical samples; MJM, XNZ, GLW, HXG, LJD, and LY performed the experiments; MJM, XNZ, GLW, HXG, LJD, XMC, and LY collected epidemiological and clinical data; SBZ helped the cell type annotation. LYT and YY performed bioinformatic analyses; and MJM and LYT drafted the manuscript. All authors reviewed and approved the final manuscript.

## Competing interesting

The authors declare no competing interests.

## METHODS

### Ethics statement

The study was conducted following the Declaration of Helsinki, and the Institutional Review Board of the Academy of Military Medical Sciences approved the study protocol (IRB number: AF/SC-08/02.46). All patients or their surrogates provided written informed consent.

### Patients

Ten patients diagnosed with SARS-CoV-2 infection were enrolled from the Fifth Hospital of Shijiazhuang from March to April 2020. SARS-CoV-2 RNA was detected in the patient’s nasopharyngeal swab or sputum specimens by real-time reverse-transcriptase PCR (RT-PCR) using the SARS-CoV-2 nucleic acid detection kit (Cat No. DA0930-DA0932, DAAN GENE Ltd., Guangzhou, China). Peripheral blood was collected from all patients during hospitalization, and blood draws from patients occurred in concert with usual care and patient’s willingness to avoid frequent blood sampling and unnecessary personal protective equipment usage. The patients’ demographic, clinical features, laboratory findings, and chest radiographs were collected from their electronic medical records.

The disease severity was defined as asymptomatic, moderate, and severe, according to the diagnostic and treatment guidelines for SARS-CoV-2 issued by the Chinese National Health Committee (Trail Version 7). Asymptomatic infection was defined as an individual who had a positive SARS-CoV-2 by RT-PCR but without any associated clinical symptoms in the preceding 14 days and during hospitalization. Moderate was defined according to the following criteria: (i) fever and respiratory symptoms; (ii) radiological signs of pneumonia. Severe was defined if satisfying at least one of the following items: (i) breathing rate ≥30/min; (ii) pulse oximeter oxygen saturation (SpO_2_) ≤93% at rest; (iii) ratio of the partial pressure of arterial oxygen (PaO_2_) to a fraction of inspired oxygen (FiO_2_) ≤300 mm Hg (1 mm Hg=0.133 kPa).

### Isolation of PMBCs

Peripheral blood mononuclear cells (PBMCs) were isolated from whole blood using density gradient centrifugation with Lymphoprep in SepMate tubes (Stemcell Technologies) in a biosafety level 2^+^ facility according to the manufactory’s instruction. Briefly, the blood was centrifuged at 1,200 × g for 10 min. PBMCs were harvested and washed twice with PBS at 400 × g for 10 min. Isolated PBMCs were frozen in cell recovery Media containing 10% DMSO (GIBCO), supplemented with 90% heat-inactivated fetal bovine serum and stored liquid nitrogen before assays analyses.

### The droplet-based single-cell RNA sequencing

Single-cell suspensions at a density of 1000 cells/μl in PBS plus 0.04% bovine serum albumin (BSA) were prepared for scRNA-seq using the Chromium Single Cell 5’ Reagent version 2 kit and Chromium Controller (10× Genomics, Pleasanton, CA), aiming for an estimated 4,000 cells per library following the manufacturer’s instructions. Briefly, 9,000 cells per reaction were loaded for gel bead-in-emulsion (GEM) generation and barcoding. GEM-RT, post-GEM-RT cleanup, and cDNA amplification were performed to isolate and amplify cDNA for library construction. Libraries were constructed using the Chromium Single Cell 5’ Reagent kit (10×Genomics) and Gel Bead Kit, Single Cell V(D)J Enrichment Kit, Human T Cell (1000005) and a Single Cell V(D)J Enrichment Kit, Human B Cell (1000016) according to the manufacturer’s protocol. Library quality and concentration were assessed according to the manufacturer’s instructions. Libraries were sequenced on an Illumina PE150.

### Single-cell RNA-seq data processing

Reads from each sample were processed with Cell Ranger (3.0.1) separately. Human reference genome GRCh38 and genome of SARS-CoV-2 were merged with corresponding GTF files used to annotate genes. The filtered matrices were then delivered into R (3.6.2) for downstream analysis. In order to demultiplex samples pooled into one sequencing run, we applied Souporcell (2.0) (Heaton et al., 2020) to separate them by individuals. Next, we used the *shared_samples.py* script in Souporcell to identify individuals. The script uses vcf files to compare shared variations when there were overlapped patients between the two runs and identifies the shared patient.

Quality control was performed using R's scater (1.14.6) (McCarthy et al., 2017) to remove cells with: (1) more than three median absolute deviations (MADs) of the log10 read counts below the median; (2) more than 3 MADs of the log10 genes detected below the median; and (3) more than 3 MADs of the genes coming from mitochondria above the median. Size factors were then considered for calculating average counts per features, and features with average counts above 0 were kept. COVID-19 genes were removed in this step as they are not detected in the data. Afterward, we used Seurat (3.2.0) (Satija et al., 2015) for data normalization and to identify highly variable genes.

### Data integration and clustering

*RunFastMNN* (Haghverdi et al., 2018) wrapped in Seurat was performed using the top 2000 highly variable genes to integrate data sets from each sample. The first 30 MNN dimension reductions were applied to construct a SNN graph and *FindClusters* with Louvain algorithms using standard Seurat pipeline. UMAP was also generated with the first 30 MNN dimension reductions to embed the data sets into two dimensions for visualization. Doublets labeled by Souporcell and clusters enriched for doublets (>15%) were removed from further analysis. We removed ribosomal and mitochondrial genes for further explore the subtypes of T/NK cells and B cells, then performed the integration and clustering again using the same strategies.

### Cell-type annotation

To annotate each cluster, we used *FindAllMarkers* in Seurat to find marker genes for each cluster and selected immune cells marker genes, as shown in the dot plot in Figure1, Figure3, and Figure5. SingleR (1.0.5) (Aran et al., 2019)was also applied to help interpretation with Monaco Immune Data (Monaco et al., 2019) (Monaco Immune Cell Data (GSE107011)) used as reference data to annotate the Th2-like cell types, which was shown in FigureS 5a. We applied *FindMarkers* in Seurat to compare the innate immune subsets that have distinct distribution in asymptomatic conditions. The selected marker genes from the comparison were shown in Figure 3d. We searched the expression of ACE2 and TNFRSF19 using the human lung atlas visualization tool (https://cellxgene.cziscience.com/d/krasnow_lab_human_lung_cell_atlas_10x-1.cxg/), and the results were shown in Supplementary Figure 5c.

The cell type abundance in Figure 2 was calculated by dividing the number of cells in a certain cell type to total cell number for a given sample, while the cell type abundance in Figure 3 and Figure 5 was calculated by dividing the number of cells in a certain cell type to the number T/NK cells and the number of B cells correspondingly. The heatmaps that present the proportions of each cell type were generated using the proportions calculated as detailed above and are scaled by row to be plotted on the same color scale. We used the Wilcoxon signed-rank test to compare the proportions of cell types between different conditions and stages. P-values were added to the plot by *stat_compare_means* function in ggpubr (0.3.0) package.

### DEG analysis

For subtypes of T cells, NK cells, and B cells with >200 cells, we aggregated the counts for cells in each sample and generated pseudobulk samples, following the data analysis workflow specified by a previous study (Crowell et al., 2020). Non-protein-coding genes and genes related to sex were removed in the counts before being analyzed by edgeR (3.28.1) (Robinson et al., 2010). *glmQLFit* and *glmQLFTest* from edgeR were used to find marker genes between each condition. Genes with a log fold change above 1 and FDR (Benjamini-Hochberg) less than 0.05 were selected. Then, genes with logCPM above 5 were shown in heatmaps in FigureS 6. GO analysis was conducted on upregulated genes using topGO (3.28.1) (Alexa et al., 2006), and the top 10 enriched GO terms were plotted.

### TCR and BCR analysis

Raw fastq files were processed with CellRanger (3.0.1) pipeline with default settings with the aforementioned reference. For TCR analysis, only cells with at least one TCR alpha chain (TRA) and one TCR beta chain (TRB) were considered as detected TCR. Moreover, each unique TRA-TRB pair was defined as a clone type. The following analyses were based on cells with detected TCR. To analyses the clonal abundance and diversity of cells from each stage, we use the alakazam (1.0.2) (Gupta et al., 2015) package. For the clonal abundance curve, the 95% confidence interval was estimated via bootstrapping. For the diversity curve, special cases of the generalized diversity index correspond to the most popular diversity measures in ecology. At q=0 different clones weight equally, regardless of sample size. As the parameter q increase from 0 to +∞, the diversity index (D) depends less on rare clones and more on common (abundant) ones. For BCR analysis, only cells with at least one heavy chain (IGH) and one light chain (IGK or IGL) were considered high-quality BCR and kept for further analysis. Furthermore, each unique IGH-IGK/IGL pair was defined as a clone type. A clone type was considered clonal if it is detected in more than one cell. All plots were generated using ggplot2 (3.3.1) (Wickham, 2016), and heatmaps were generated using pheatmap (1.0.12) unless otherwise specified.

### Data and Code Availability

Raw and processed data are available on CNGB Nucleotide Sequence Archive (CNSA: https://db.cngb.org/cnsa) with accession number CNP0001250. The codes supporting the current study are available from Github (https://github.com/YOU-k/covid_analysis).

## REFERENCES

Abel, A.M., Yang, C., Thakar, M.S., and Malarkannan, S. (2018). Natural Killer Cells: Development, Maturation, and Clinical Utilization. Front Immunol 9, 1869.

Alexa, A., Rahnenfuhrer, J., and Lengauer, T. (2006). Improved scoring of functional groups from gene expression data by decorrelating GO graph structure. Bioinformatics 22, 1600–1607.

Aran, D., Looney, A.P., Liu, L., Wu, E., Fong, V., Hsu, A., Chak, S., Naikawadi, R.P., Wolters, P.J., Abate, A.R., et al. (2019). Reference-based analysis of lung single-cell sequencing reveals a transitional profibrotic macrophage. Nat Immunol 20, 163–172.

Chen, G., Wu, D., Guo, W., Cao, Y., Huang, D., Wang, H., Wang, T., Zhang, X., Chen, H., Yu, H., et al. (2020a). Clinical and immunological features of severe and moderate coronavirus disease 2019. The Journal of clinical investigation 130, 2620–2629.

Chen, N., Zhou, M., Dong, X., Qu, J., Gong, F., Han, Y., Qiu, Y., Wang, J., Liu, Y., Wei, Y., et al. (2020b). Epidemiological and clinical characteristics of 99 cases of 2019 novel coronavirus pneumonia in Wuhan, China: a descriptive study. Lancet (London, England) 395, 507–513.

Crowell, H.L., Soneson, C., Germain, P.-L., Calini, D., Collin, L., Raposo, C., Malhotra, D., and Robinson, M.D. (2020). On the discovery of subpopulation-specific state transitions from multi-sample multi-condition single-cell RNA sequencing data. bioRxiv, 713412.

Fregni, G., Maresca, A.F., Jalbert, V., Caignard, A., Scott-Algara, D., Cramer, E.B., Rouveix, E., Bene, M.C., and Capron, C. (2013). High number of CD56(bright) NK-cells and persistently low CD4+ T-cells in a hemophiliac HIV/HCV co-infected patient without opportunistic infections. Virol J 10, 33.

Gandhi, M., Yokoe, D.S., and Havlir, D.V. (2020). Asymptomatic Transmission, the Achilles' Heel of Current Strategies to Control Covid-19. N Engl J Med 382, 2158–2160.

Giamarellos-Bourboulis, E.J., Netea, M.G., Rovina, N., Akinosoglou, K., Antoniadou, A., Antonakos, N., Damoraki, G., Gkavogianni, T., Adami, M.E., Katsaounou, P., et al. (2020). Complex Immune Dysregulation in COVID-19 Patients with Severe Respiratory Failure. Cell Host Microbe 27, 992–1000 e1003.

Gibbons, D.L., Abeler-Dorner, L., Raine, T., Hwang, I.Y., Jandke, A., Wencker, M., Deban, L., Rudd, C.E., Irving, P.M., Kehrl, J.H., et al. (2011). Cutting Edge: Regulator of G protein signaling-1 selectively regulates gut T cell trafficking and colitic potential. J Immunol 187, 2067–2071.

Grifoni, A., Weiskopf, D., Ramirez, S.I., Mateus, J., Dan, J.M., Moderbacher, C.R., Rawlings, S.A., Sutherland, A., Premkumar, L., Jadi, R.S., et al. (2020). Targets of T Cell Responses to SARS-CoV-2 Coronavirus in Humans with COVID-19 Disease and Unexposed Individuals. Cell 181, 1489–1501 e1415.

Gupta, N.T., Vander Heiden, J.A., Uduman, M., Gadala-Maria, D., Yaari, G., and Kleinstein, S.H. (2015). Change-O: a toolkit for analyzing large-scale B cell immunoglobulin repertoire sequencing data. Bioinformatics 31, 3356–3358.

Haghverdi, L., Lun, A.T.L., Morgan, M.D., and Marioni, J.C. (2018). Batch effects in single-cell RNA-sequencing data are corrected by matching mutual nearest neighbors. Nat Biotechnol 36, 421–427.

Heaton, H., Talman, A.M., Knights, A., Imaz, M., Gaffney, D.J., Durbin, R., Hemberg, M., and Lawniczak, M.K.N. (2020). Souporcell: robust clustering of single-cell RNA-seq data by genotype without reference genotypes. Nat Methods 17, 615–620.

Herman, J.S., Sagar, and Grun, D. (2018). FateID infers cell fate bias in multipotent progenitors from single-cell RNA-seq data. Nat Methods 15, 379–386.

Huang, C., Wang, Y., Li, X., Ren, L., Zhao, J., Hu, Y., Zhang, L., Fan, G., Xu, J., Gu, X., et al. (2020). Clinical features of patients infected with 2019 novel coronavirus in Wuhan, China. Lancet (London, England) 395, 497–506.

Jose, R.J., and Manuel, A. (2020). COVID-19 cytokine storm: the interplay between inflammation and coagulation. The Lancet Respiratory medicine 8, e46–e47.

Kojima, T., Morikawa, Y., Copeland, N.G., Gilbert, D.J., Jenkins, N.A., Senba, E., and Kitamura, T. (2000). TROY, a newly identified member of the tumor necrosis factor receptor superfamily, exhibits a homology with Edar and is expressed in embryonic skin and hair follicles. J Biol Chem 275, 20742–20747.

Krausgruber, T., Fortelny, N., Fife-Gernedl, V., Senekowitsch, M., Schuster, L.C., Lercher, A., Nemc, A., Schmidl, C., Rendeiro, A.F., Bergthaler, A., et al. (2020). Structural cells are key regulators of organ-specific immune responses. Nature 583, 296–302.

Kumar, B.V., Ma, W., Miron, M., Granot, T., Guyer, R.S., Carpenter, D.J., Senda, T., Sun, X., Ho, S.H., Lerner, H., et al. (2017). Human Tissue-Resident Memory T Cells Are Defined by Core Transcriptional and Functional Signatures in Lymphoid and Mucosal Sites. Cell Rep 20, 2921–2934.

Lee, J.S., Park, S., Jeong, H.W., Ahn, J.Y., Choi, S.J., Lee, H., Choi, B., Nam, S.K., Sa, M., Kwon, J.S., et al. (2020). Immunophenotyping of COVID-19 and influenza highlights the role of type I interferons in development of severe COVID-19. Science immunology 5.

Liao, M., Liu, Y., Yuan, J., Wen, Y., Xu, G., Zhao, J., Cheng, L., Li, J., Wang, X., Wang, F., et al. (2020). Single-cell landscape of bronchoalveolar immune cells in patients with COVID-19. Nature medicine 26, 842–844.

Long, Q.X., Tang, X.J., Shi, Q.L., Li, Q., Deng, H.J., Yuan, J., Hu, J.L., Xu, W., Zhang, Y., Lv, F.J., et al. (2020). Clinical and immunological assessment of asymptomatic SARS-CoV-2 infections. Nature medicine.

Lucas, C., Wong, P., Klein, J., Castro, T.B.R., Silva, J., Sundaram, M., Ellingson, M.K., Mao, T., Oh, J.E., Israelow, B., et al. (2020). Longitudinal analyses reveal immunological misfiring in severe COVID-19. Nature.

Mathew, D., Giles, J.R., Baxter, A.E., Oldridge, D.A., Greenplate, A.R., Wu, J.E., Alanio, C., Kuri-Cervantes, L., Pampena, M.B., D'Andrea, K., et al. (2020). Deep immune profiling of COVID-19 patients reveals distinct immunotypes with therapeutic implications. Science.

McCarthy, D.J., Campbell, K.R., Lun, A.T., and Wills, Q.F. (2017). Scater: pre-processing, quality control, normalization and visualization of single-cell RNA-seq data in R. Bioinformatics 33, 1179–1186.

Mehta, P., McAuley, D.F., Brown, M., Sanchez, E., Tattersall, R.S., and Manson, J.J. (2020). COVID-19: consider cytokine storm syndromes and immunosuppression. Lancet (London, England) 395, 1033–1034.

Michel, T., Poli, A., Cuapio, A., Briquemont, B., Iserentant, G., Ollert, M., and Zimmer, J. (2016). Human CD56bright NK Cells: An Update. J Immunol 196, 2923–2931.

Monaco, G., Lee, B., Xu, W., Mustafah, S., Hwang, Y.Y., Carre, C., Burdin, N., Visan, L., Ceccarelli, M., Poidinger, M., et al. (2019). RNA-Seq Signatures Normalized by mRNA Abundance Allow Absolute Deconvolution of Human Immune Cell Types. Cell Rep 26, 1627–1640 e1627.

World Health Organization. (2020). Coronavirus disease (COVID-19) Situation Report – 177.

Raoult, D., Zumla, A., Locatelli, F., Ippolito, G., and Kroemer, G. (2020). Coronavirus infections: Epidemiological, clinical and immunological features and hypotheses. Cell stress 4, 66–75.

Robinson, M.D., McCarthy, D.J., and Smyth, G.K. (2010). edgeR: a Bioconductor package for differential expression analysis of digital gene expression data. Bioinformatics 26, 139–140.

Satija, R., Farrell, J.A., Gennert, D., Schier, A.F., and Regev, A. (2015). Spatial reconstruction of single-cell gene expression data. Nat Biotechnol 33, 495–502.

Scharenberg, M., Vangeti, S., Kekalainen, E., Bergman, P., Al-Ameri, M., Johansson, N., Sonden, K., Falck-Jones, S., Farnert, A., Ljunggren, H.G., et al. (2019). Influenza A Virus Infection Induces Hyperresponsiveness in Human Lung Tissue-Resident and Peripheral Blood NK Cells. Front Immunol 10, 1116.

Schulte-Schrepping, J., Reusch, N., Paclik, D., Bassler, K., Schlickeiser, S., Zhang, B., Kramer, B., Krammer, T., Brumhard, S., Bonaguro, L., et al. (2020). Severe COVID-19 Is Marked by a Dysregulated Myeloid Cell Compartment. Cell.

Skarica, M., Eckstein, C., Whartenby, K.A., and Calabresi, P.A. (2011). Novel mechanisms of immune modulation of natalizumab in multiple sclerosis patients. J Neuroimmunol 235, 70–76.

Sungnak, W., Huang, N., Becavin, C., Berg, M., Queen, R., Litvinukova, M., Talavera-Lopez, C., Maatz, H., Reichart, D., Sampaziotis, F., et al. (2020). SARS-CoV-2 entry factors are highly expressed in nasal epithelial cells together with innate immune genes. Nature medicine 26, 681–687.

Swain, S.L., McKinstry, K.K., and Strutt, T.M. (2012). Expanding roles for CD4(+) T cells in immunity to viruses. Nat Rev Immunol 12, 136–148.

Travaglini, K.J., Nabhan, A.N., Penland, L., Sinha, R., Gillich, A., Sit, R.V., Chang, S., Conley, S.D., Mori, Y., Seita, J., et al. (2020). A molecular cell atlas of the human lung from single cell RNA sequencing. bioRxiv, 742320.

Wickham, H. (2016). ggplot2: Elegant Graphics for Data Analysis (Springer International Publishing).

Wilk, A.J., Rustagi, A., Zhao, N.Q., Roque, J., Martínez-Colón, G.J., McKechnie, J.L., Ivison, G.T., Ranganath, T., Vergara, R., Hollis, T., et al. (2020). A single-cell atlas of the peripheral immune response in patients with severe COVID-19. Nature medicine 26, 1070–1076.

Zhang, J.Y., Wang, X.M., Xing, X., Xu, Z., Zhang, C., Song, J.W., Fan, X., Xia, P., Fu, J.L., Wang, S.Y., et al. (2020). Single-cell landscape of immunological responses in patients with COVID-19. Nat Immunol.

Zhou, Z., Ren, L., Zhang, L., Zhong, J., Xiao, Y., Jia, Z., Guo, L., Yang, J., Wang, C., Jiang, S., et al. (2020). Heightened Innate Immune Responses in the Respiratory Tract of COVID-19 Patients. Cell Host Microbe 27, 883–890 e882.

Zhu, L., Yang, P., Zhao, Y., Zhuang, Z., Wang, Z., Song, R., Zhang, J., Liu, C., Gao, Q., Xu, Q., et al. (2020). Single-Cell Sequencing of Peripheral Mononuclear Cells Reveals Distinct Immune Response Landscapes of COVID-19 and Influenza Patients. Immunity.

